# GRP78 promotes stemness in normal and neoplastic cells

**DOI:** 10.1101/720698

**Authors:** Clay Conner, Tyson W. Lager, Ian H. Guldner, Min-Zu Wu, Yuriko Hishida, Tomoaki Hishida, Sergio Ruiz, Amanda E. Yamasaki, Robert C. Gilson, Juan Carlos Izpisua Belmonte, Peter C. Gray, Jonathan A. Kelber, Siyuan Zhang, Athanasia D. Panopoulos

## Abstract

Reliable approaches to identify stem cell mechanisms that mediate aggressive cancer could have great therapeutic value, based on the growing evidence of embryonic signatures in metastatic cancers. However, how to best identify and target stem-like mechanisms aberrantly acquired by cancer cells has been challenging. We harnessed the power of reprogramming to examine GRP78, a chaperone protein generally restricted to the endoplasmic reticulum in normal tissues, but which is expressed on the cell surface of human embryonic stem cells and many cancer types. We have discovered that (1) cell surface GRP78 (sGRP78) is expressed on iPSCs and is important in reprogramming, (2) sGRP78 promotes cellular functions in both pluripotent and breast cancer cells (3) overexpression of GRP78 in breast cancer cells leads to an induction of a CD24^-^/CD44^+^ tumor initiating cell (TIC) population (4) sGRP78^+^ breast cancer cells are enriched for stemness genes and appear to be a subset of TICs (5) sGRP78^+^ breast cancer cells show an enhanced ability to seed metastatic organ sites *in vivo*. These collective findings show that GRP78 has important functions in regulating both pluripotency and oncogenesis, and suggest that sGRP78 marks a stem-like population in breast cancer cells that has increased metastatic potential *in vivo*.

## INTRODUCTION

Accumulating evidence has shown the presence of embryonic stem-cell programs in cancer cells that contribute to aggressive malignancy [1-3]. Previous work has also demonstrated that common pathways critical in oncogenesis parallel many pathways important in the induction of pluripotency [4-6]. Recently, a group using machine learning found an increased stemness index in metastatic tumors across many cancer types [7]. A study using single-cell analysis specifically correlated a stem-cell program to human metastatic breast cancer cells, expanding on the existing paradigm that pairs stem cell programs with disease aggressiveness [8]. These findings support the concept that an embryonic program may be aberrantly retained and/or reactivated to be exploited in cancer. Thus, examining pluripotent stem cells in relation to cancer could provide a powerful approach to gaining insight into critical mechanisms regulating aggressive tumors.

Glucose-regulated protein 78 (GRP78; also known as heat shock 70 kDA protein 5, HSPA5) is a stress inducible endoplasmic reticulum (ER) chaperone protein that is part of the larger heat shock protein superfamily. GRP78 is typically localized in the ER to assist in the protein folding and assembly of membrane or secreted proteins [9]. GRP78 overexpression has been observed clinically for many different cancer types, where it has been shown to correlate to metastasis and poor patient outcome [10, 11]. These findings are supported by genetically engineered mouse models and tissue culture systems, which have also shown roles for the overexpression of GRP78 in malignancy [12], invasive and aggressive phenotypes [13, 14], metastasis [15], and drug resistant properties [16]. Previous reports indicate that GRP78 is aberrantly localized to the cell surface in many types of cancer (e.g. breast, pancreas, lung, ovarian, colon, melanoma) [11, 17], where it has been linked to the regulation of critical signaling pathways [18-22].

We have previously reported that GRP78 is expressed on the surface of human embryonic stem cells [20]. Furthermore, Spike et. al. found functions for cell surface GRP78 in fetal and adult mammary stem/progenitor populations [22]. These collective findings suggest that aberrant cancer functions of GRP78 may be revealing an embryonic function of GRP78 that is inappropriately reactivated and exploited, or that adult stem cells could be retaining certain embryonic mechanisms of GRP78 that then become aberrant. In this study, we examined the functions of GRP78 in breast cancer and in the acquisition and maintenance of pluripotency, which revealed important insights into understanding how cancer cells acquire and/or exploit embryonic stem cell mechanisms.

## RESULTS

To examine the role of GRP78 in regulating the acquisition of an embryonic stem cell state, we first modified GRP78 expression levels during reprogramming. Knockdown of GRP78 (Supplementary Figure S1) during reprogramming of human fibroblasts following the transduction of *OCT4, SOX2, KLF4* and *cMYC* (referred to as 4F) led to a significant decrease in the number of iPSC colonies present, as judged by expression of the pluripotent marker NANOG in the resulting colonies (Figure 1A). Conversely, overexpression of GRP78 (Supplementary Figure S1) in conjunction with 4F led to a significant induction of iPSC colonies (∼2.5 fold), and in the absence of *cMYC* (referred to as 3F) during reprogramming, an even greater increase (∼4-fold) compared to controls (Figure 1B, 1C). When fibroblasts were ‘primed’ with overexpression of GRP78 two days prior to being transduced with the 4F to induce reprogramming, the increase was 6-fold (Figure 1D). To examine GRP78 expression after reprogramming, we stained induced pluripotent stem cells (iPSCs) with GRP78 and the cell membrane protein E-cadherin [23]. Our results show that GRP78 is co-expressed with E-cadherin on the cell surface of iPSCs, in agreement with our previous findings of GRP78 on the cell surface of human embryonic stem cells [20] (Figure 1E). These results indicate that GRP78 plays an important role in the reprogramming process, and that GRP78 is expressed on the cell surface of iPSCs.

**Figure 1:**
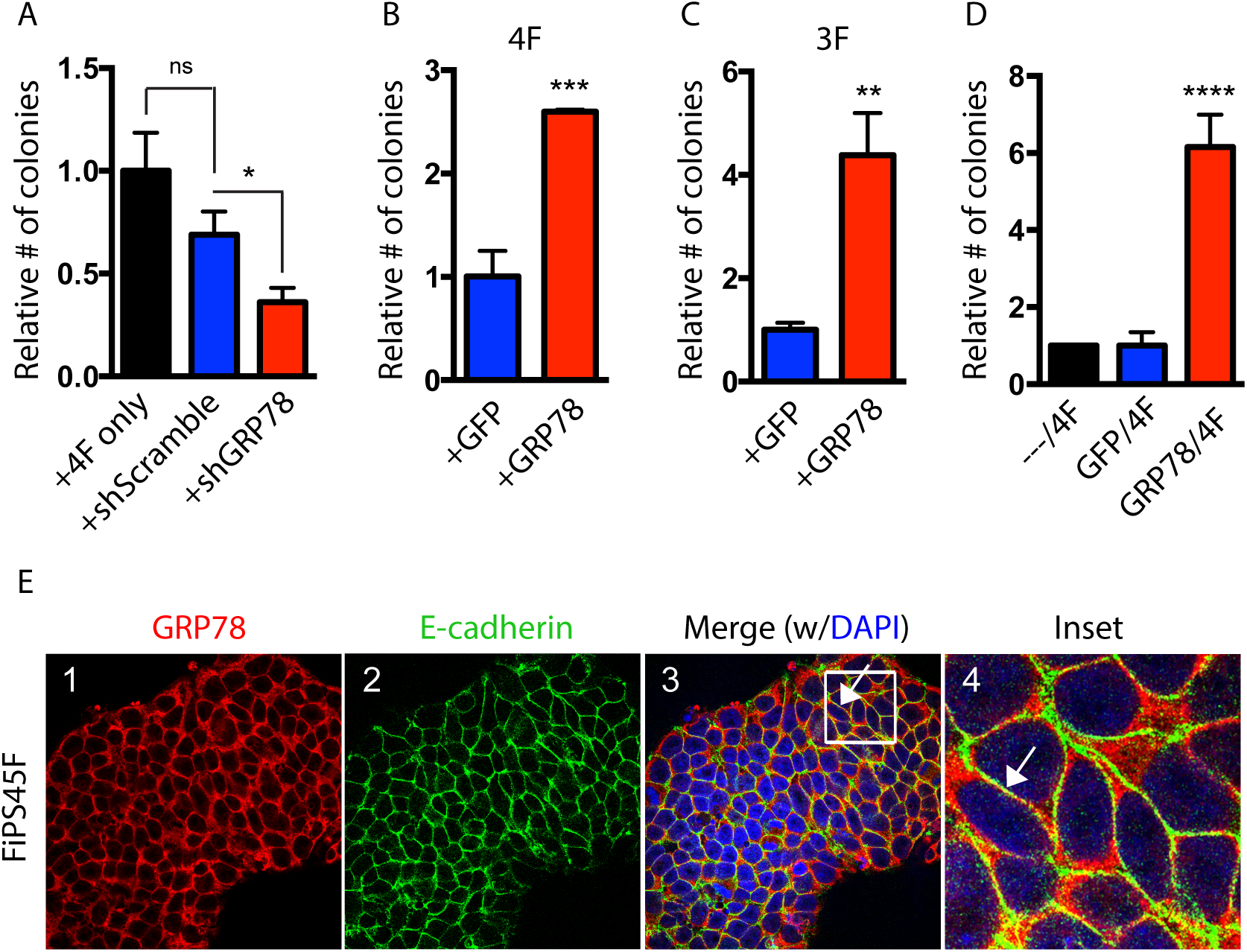
GRP78 is important for reprogramming and is expressed on the surface of iPSCs. (A) Human keratinocytes were retrovirally infected with *OCT4, SOX2, KLF4* and *c-MYC* (OSKM) either alone (4F) or in the presence of shGRP78 or shScramble control. Resulting colonies (∼20 days after infection) were stained for Nanog and colony numbers determined relative to the control. (B) Keratinocytes were retrovirally infected with OSKM (4F) or (C) with OSK (3F), in addition to a retrovirus expressing GRP78 or a GFP control. The number of Nanog positive colonies are shown relative to the control. (D) Keratinocytes were retrovirally infected with 4F following a 2-day prime with either a retrovirus expressing GRP78 or a GFP control. Nanog positive colony numbers are shown relative to the control. (E) A previously characterized human iPSC line derived from fibroblasts (FiPS4F5 [40]) was examined by immunofluorescence for GRP78 (red), and E-cadherin (green, a cell surface marker), and both markers were found to colocalize together (Merge, Inset; arrows). 4,6-Diamidino-2-phenylindole (DAPI) staining shows nuclei. Results were quantified from triplicate samples, and are representative of at least three independent experiments. Error bars depict the standard error mean (SEM). \**p*<0.05; \*\**p*<0.01; \*\*\**p*<0.001; \*\*\*\**p*<0.0001; ns = not significant.

Cell surface GRP78 (referred to herein as sGRP78) on cancer cells has already emerged as a potential chemotherapeutic target [11], but the potential cell surface function of sGRP78 in human pluripotent stem cells remained unknown. We found that GRP78 relocalization appears by the 1-2 cell stage of reprogramming (Figure 2A), suggesting that the presence of sGRP78 is an early event in the acquisition of pluripotency. When using a GRP78 antibody, that has previously been reported to disrupt sGRP78 function [20], or matched IgG control at different stages during reprogramming, the efficiency of reprogramming was inhibited (Figures 2B and 2C). The kinetics of antibody treatment and the effects on colony number suggested that sGRP78 functioned during the reprogramming process (Figure 2C). GRP78 antibody treatment at later stages, when colonies were already present, did not statistically affect colony number (Figure 2C). However, if the inhibitory GRP78 antibody treatment at later stages (i.e. after colony development) was causing a reduction in colony size (i.e. causing some of the cells to die and/or differentiate), this would not be indicated by only measuring colony number. Therefore, to next examine the function of GRP78 on the cell surface of human pluripotent stem cells (PSCs), we treated PSCs that express GFP under the control of an OCT4 promoter [24] with the inhibitory GRP78 antibody. Treatment of PSCs with this antibody decreased PSC number, but did not impact pluripotency of the remaining population, as judged by OCT4-GFP expression levels (Figure 2D). Thus, the loss of PSCs from inhibiting sGRP78 was not due to differentiation, and instead was likely due to survival and/or proliferation mechanisms. Cell cycle analysis of PSCs following GRP78 antibody (or IgG control) treatments demonstrated that the cell cycle was not changing (Figure 2E). Thus, coupled with the lack of differentiation, this suggests that sGRP78 is important in maintaining PSC cell survival. Interestingly, inducing GRP78 overexpressing in the human basal (i.e. ER-/PR-/Her2-) breast cancer cell line MDA-MB-231 caused an increased resistance to the chemotherapeutic drug cisplatin (Figure 2F). The ability of GRP78 to regulate breast cancer cell survival was also cell surface dependent, since inhibiting cell surface function of GRP78 with the inhibitory antibody to GRP78 resulted in a higher susceptibility to cisplatin treatment (Figure 2F). Importantly, this resistance or changed susceptibility with GRP78 antibody was not seen in the absence of GRP78 overexpression, or when inducing overexpressing of a GFP control (Supplementary Figures S2 and S3).

**Figure 2:**
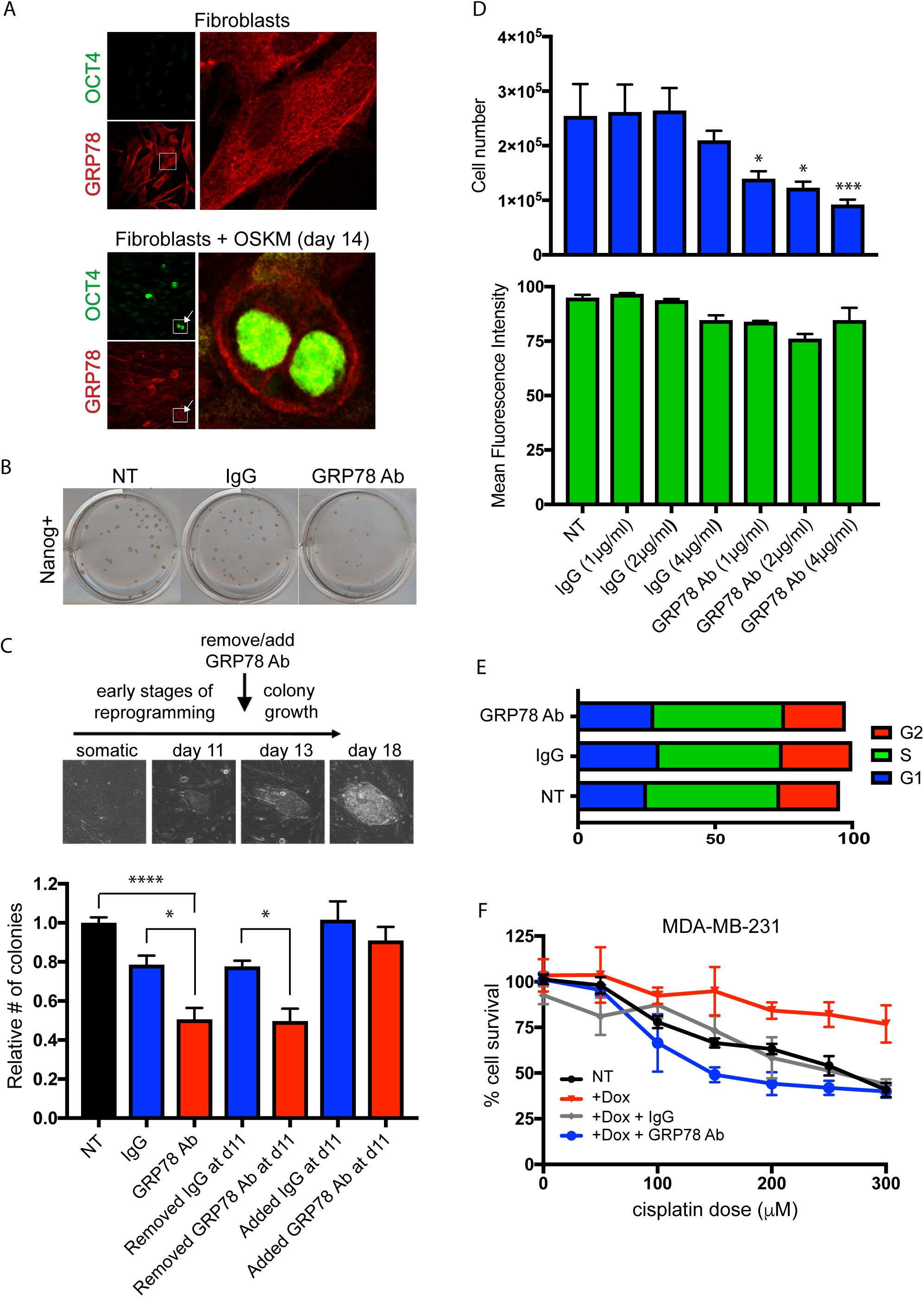
Cell surface GRP78 plays a significant role in reprogramming and pluripotent stem cell and breast cancer cell function. (A) Fibroblasts (dFib-OCT4^GFP^ cells [29]) were plated on coverslips and infected with *OCT4, SOX2, KLF4* and *c-MYC* (OSKM) to initiate reprogramming. At 14 days post infection, cells were fixed and stained with OCT4 (green) and GRP78 (red). Day 14 shows the appearance of OCT4-positive cells (arrows, inset) at this timepoint, indicative of cells that have undergone reprogramming back to a pluripotent state. Note the change in GRP78 localization, primarily at the cell surface (insets). Fibroblasts that were not infected with virus containing OSKM were used as a negative control. (B-C) Keratinocytes were infected with OSKM, plated, and cells were treated with media only, a GRP78 inhibiting antibody or IgG control throughout, or for the indicated timepoints. Following approximately 20 days after OSKM infection, resulting colonies were stained for Nanog (representative staining at this timepoint shown in B, used as the endpoint in the assay), and colony numbers determined relative to media only control. Note that cell surface expression of GRP78 (sGRP78) appears to be required for the reprogramming process (e.g. early anti-GRP78 timepoints affect colony numbers), rather than affecting colony number after colonies had already formed. Morphological pictures of cells throughout the reprogramming process (all at same magnification) are shown. (D-E) Pluripotent stem cells (PSCs) where GFP is driven by the OCT4 promoter [24] were plated in triplicate and treated with media only, media containing GRP78 antibody, or respective IgG control, at the concentrations indicated. Media was replaced daily for four days. Cell counts were determined by trypan blue exclusion (D, upper panel). GFP expression (as a measure of OCT4 expression) was analyzed by flow cytometry (D, lower panel). (E) Cell cycle distribution determined by propidium iodide (PI) staining followed by flow cytometry analysis. (F) MDA-MB-231 breast cancer cells that overexpressed RFP-GRP78 under a doxycycline (Dox)-inducible promoter, in the absence or presence of a GRP78 inhibiting antibody, were used in MTT-based survival assays to assess cisplatin-induced apoptosis. Note that the resistance to cisplatin-induced death that overexpression of GRP78 provides (red line) is dependent on cell surface expression of GRP78 (blue line). Results were quantified from triplicate samples, and are representative of at least three independent experiments. Error bars depict the standard error mean (SEM). \**p*<0.05; \*\**p*<0.01; \*\*\**p*<0.001; \*\*\*\**p*<0.0001; ns = not significant.

Although sGRP78 shared similar functions on stem cells and cancer cells, whether or not sGRP78 expression was marking and/or affecting a stem-like subpopulation within breast cancer remained unclear. To begin to analyze the potential role of sGRP78 expression in stemness in cancer, we performed experiments with the breast cancer cell lines MCF7 (luminal A subtype [25]) and MDA-MB-231 (basal subtype [25]) that contain low or high levels of previously defined CD24^-^/CD44^+^ tumor initiating cells (TICs) respectively, a cell population with breast cancer cells shown to be able to generate tumors with much higher efficiency in *in vivo* xenograft transplantation assays [26]. Interestingly, overexpression of GRP78 in both breast cancer cell lines caused an induction of the TIC subpopulation (i.e. CD24^-^/CD44^+^ cells) [26] (Figures 3A and 3B). Overexpressing GRP78 also induced the expression of aldehyde dehydrogenase 1 (ALDH1A), a marker associated with stemness and cancer [27] (Supplementary Figure S4).

**Figure 3:**
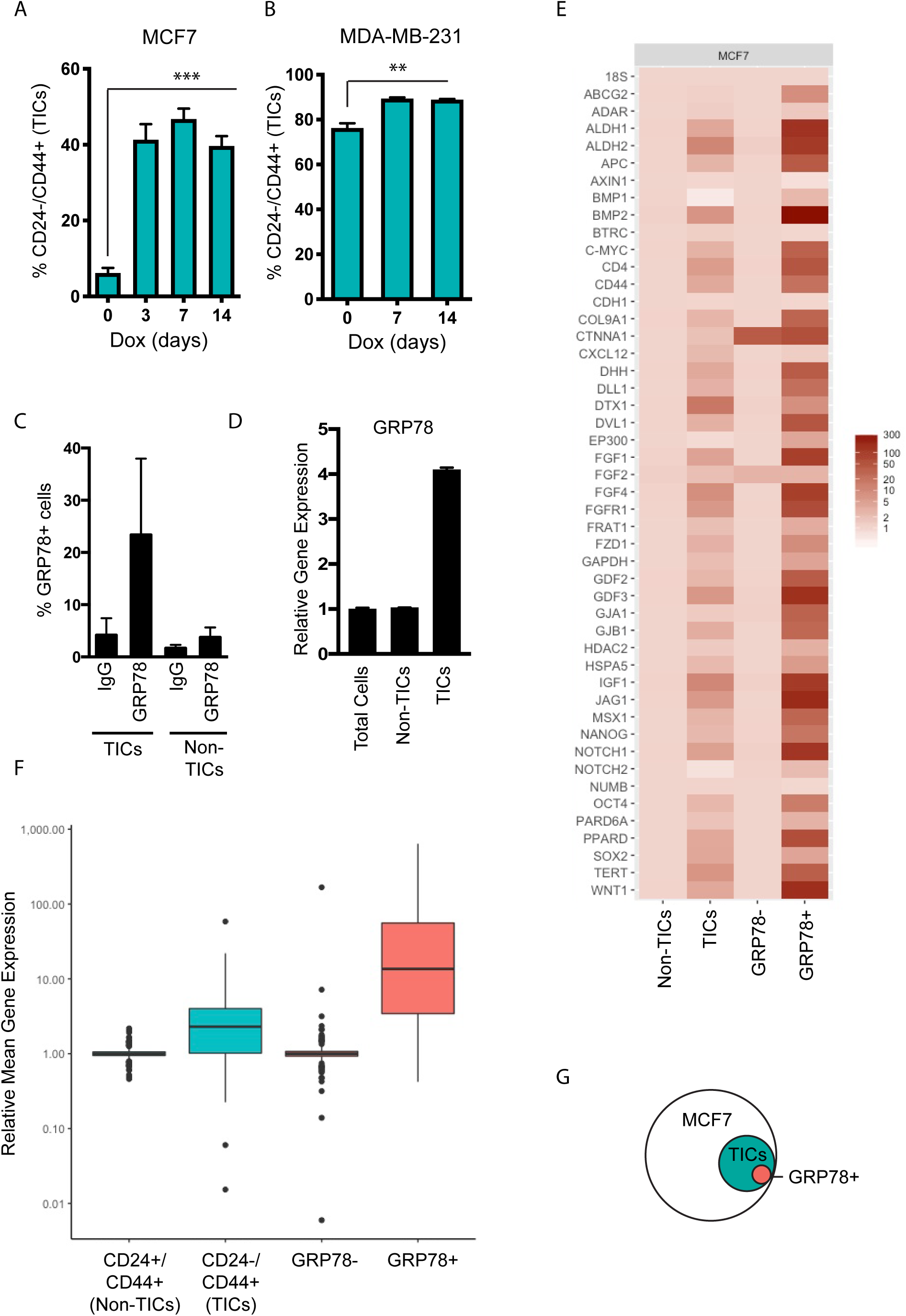
GRP78 induces tumor initiating cell (TIC) populations in breast cancer, and sGRP78^+^ cells are a subset of TICs that show elevated levels of genes important in stem cell functions. (A) MCF7-RFP-GRP78 and (B) MDA-MB-231-RFP-GRP78 cells show an increase in the CD44^+^/CD24^-^ tumor initiating population (TIC) following Dox treatment (to induce GRP78 expression) by flow cytometry. (C) Relative CT values of 46 stemness-related genes (as previously described [29]) were compared between RNA obtained from sGRP78^+/-^ and TIC/Non-TIC subpopulations isolated by FACS. (D) Relative quantification of genes shown in C demonstrating that sGRP78^+^ cells express higher levels of stemness genes compared to other populations. (E) MCF7 cells were labeled with CD44, CD24, and IgG or GRP78 and examined by flow cytometry, and show that sGRP78^+^ cells are predominantly located in the TIC (CD24^-^/CD44^+^) subpopulation compared to the Non-CSC (CD24^+^/CD44^+^) population. (F) Relative transcript levels of GRP78 from total MCF7 cells or Non-TIC or TIC sorted populations. (F) Schematic representing (E). Results are representative of at least three independent experiments. Error bars depict the standard error mean (SEM). \**p*<0.05; \*\**p*<0.01; \*\*\**p*<0.001; \*\*\*\**p*<0.0001; ns = not significant.

To next compare sGRP78 populations to the previously established TIC populations, we first examined the cell surface expression of GRP78 relative to the CD24^-^/CD44^+^ population. We focused on MCF7 cells, since they express low numbers of TICs compared to MDA-MB-231 (Figures 3A and 3B), thus enabling differences between the various populations to be better examined. Interestingly, flow cytometry analysis revealed that the sGRP78^+^ cells were a subpopulation within the TIC population (Figure 3C). Furthermore, GRP78 gene expression levels were higher in TICs (Figure 3D), in agreement with previous reports [28]. Thus, if previous studies have identified that TIC subsets mark a heterogeneous but potentially enriched stem-like population, is it possible that sGRP78 expression marks a subpopulation of ‘purer’ stem-like cells? To test this question, we sorted sGRP78^+^ cells and CD24^-^/CD44^+^ cells from MCF7 by fluorescence activated cell sorting (FACS), isolated RNA, and performed gene expression analysis of a panel of stem-cell related genes [29] (Figure 3E). Expression of stem genes were increased in the TICs compared to the Non-TIC population (CD24^+^/CD44^+^), as expected (Figures 3E and 3F). Strikingly, however, mean collective gene expression analysis demonstrated that sGRP78^+^ cells showed a higher overall expression of stem genes, even compared to the TIC population (Figure 3F). Thus, our combined results suggest that sGRP78^+^ cells contain higher levels of key genes important in stem cell functions, and are a subset of a previously established TIC population in breast cancer cells (Figure 3G).

To determine if sGRP78^+^ cells had functional relevance, we performed *in vitro* and *in vivo* assays of tumorigenic potential. TICs isolated from MCF7 cells showed an increased ability to generate colonies in anchorage-independent soft agar assays compared to Non-TIC controls, as expected (Figure 4A). The number generated from sGRP78^+^ isolated cells was even more pronounced, showing a statistically significant higher number of colonies than TICs. Furthermore, the colonies generated from sGRP78^+^ cells were bigger in size than both TICs and Non-TIC controls (Figure 4B). Since the *in vitro* assays suggested that the sGRP78^+^ population had an increased tumorigenic potential, we next examined the tumorigenicity of this population *in vivo*. TIC or sGRP78^+^ sorted cells, or total MCF7 cells (which served as a control as they contain low levels of both TIC and sGRP78^+^ populations) were fluorescently labeled, and intracardiacally injected into immunodeficient mice. The presence of fluorescent cells in known sites of breast cancer metastasis (e.g. lung, brain) was examined at both 2 days after injection (to determine the ability of each population to ‘seed’ to organ sties of metastasis), and 1 month (to determine the ability of each population to generate tumor growths at metastatic sites). We found that while total MCF7 cells and previously established TICs were able to ‘seed’ to the lung and brain within 2 days after injection, sGRP78^+^ cells showed a dramatically enhanced ability to seed the lungs and brains of mice (Figures 4C and 4D). Similarly, after 1 month, the sGRP78^+^ cells continued to show an increased ability to generate tumors in both lung and brain (Figures 4E and 4F), Furthermore, when examining the lungs of each mouse group after 1 month, the colonization architecture of sGRP78^+^ populations was substantially larger than TIC populations (Figure 4G), suggestive of a more aggressive behavior. These collective findings suggest that sGRP78 marks a stem-like population in breast cancer cells that has increased metastatic potential *in vivo*.

**Figure 4:**
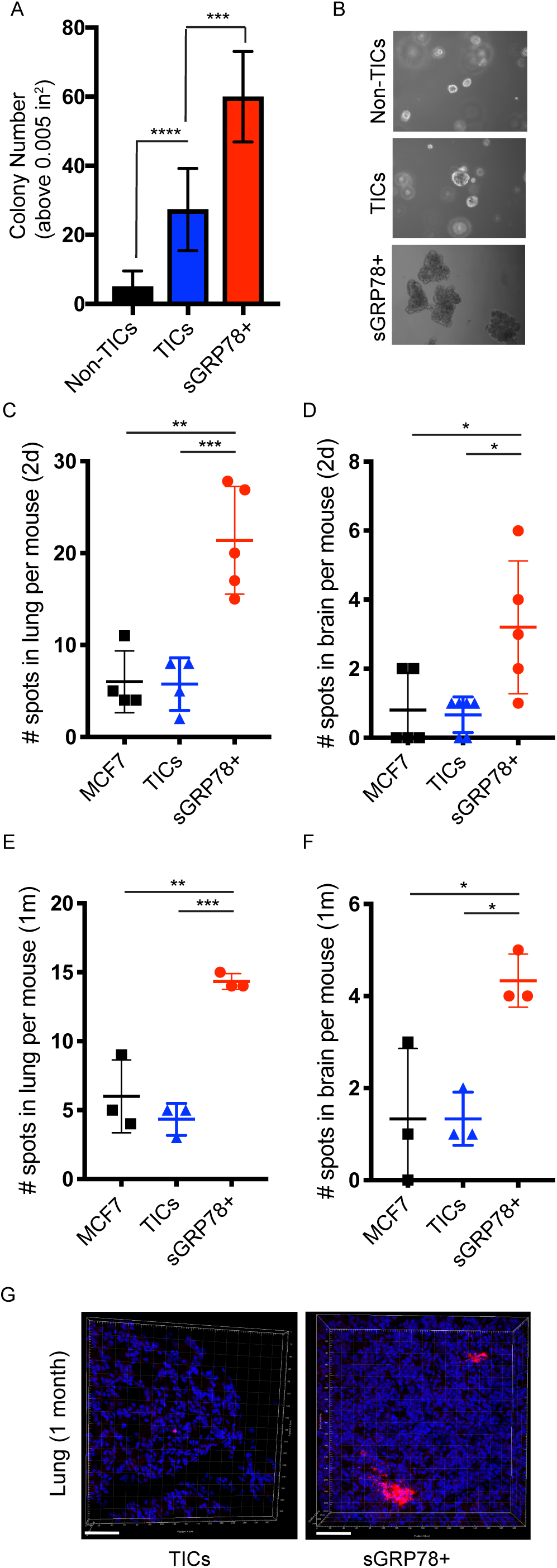
sGRP78^+^ cells show increased tumorigenesisS potential *in vitro* and *in vivo*. (A) MCF7 TICs (CD24^-^/CD44^+^), Non-TICs (CD24^+^/CD44^+^), or sGRP78^+^ cells were sorted by FACS and seeded into soft agar conditions to assess *in vitro* tumorigenicity. Cells were fed daily for 10 days. To assess live colony growth, cells were stained with INT-Violet. Total colony number (size above 0.005 in^2^) was scored using Image J Analysis. (B) Sample pictures representing (A). (C-G) Immunocompromised were injected with either unsorted DiI-stained MCF7 cells, or sorted (TIC or sGRP78^+^) DiI-stained MCF7 cells, and organs were collected 2 days later (C-D) or 1 month later (E-G) and DiI-stained cells were manually counted (C-F) or imaged (G). sGRP78^+^ cells led to significantly more breast cancer cells in lung and brain compared to other populations, both at seeding timepoints and after 1 month. (G) The size of the larger tumors made by sGRP78^+^ cells compared to TICs is shown for lung. Results are representative of at least three independent experiments. Error bars depict the standard error mean (SEM). \**p*<0.05; \*\**p*<0.01; \*\*\**p*<0.001; \*\*\*\**p*<0.0001; ns = not significant.

## DISCUSSION

GRP78 can act both as an ER-chaperone to assist in protein folding, and at the cell surface where it has been linked to the regulation of critical signaling pathways. The ER-chaperone function of GRP78 is ubiquitous, but many studies have reported elevated levels of GRP78 in various cancers [11]. The expression of GRP78 at the cell surface (i.e. sGRP78) has been restricted *in vivo* to cancer cells [11], and stem cells - specifically a subset of hematopoietic stem cells [30], fetal and mammary stem cells [22], and human embryonic stem cells [20]. Thus, we sought to investigate, are the “aberrant” functions of GRP78 reported in cancer actually the repurposing of stem cell functions of GRP78? Here we report that GRP78 is important in somatic cell reprogramming, in pluripotent stem cell and breast cancer cell functions, and in promoting tumor initiating cell populations within breast cancer. Furthermore, our data also shows that sGRP78 marks a subpopulation within breast cancer that has elevated expression of genes important in stem cell functions, and increased tumorigenicity *in vitro* and *in vivo*. These collective findings support the concept that the aberrant cancer functions of GRP78 could represent embryonic functions of GRP78 that have been inappropriately retained/reacquired and exploited in cancer cells to help facilitate their aggressiveness.

We first examined GRP78 in somatic cell reprogramming. Inhibiting total GRP78 levels in somatic cells during reprogramming reduced reprogramming efficiency, in support of the important role for GRP78 in early embryogenesis [31]. Conversely, overexpressing GRP78 levels during reprogramming increased reprogramming efficiency. It is of interest that elevated levels of GRP78 can contribute to an increase in reprogramming, especially in light of the increase in GRP78 levels that have been reported for many tumor types [11]. When paired with the fact that increasing GRP78 levels led to an increase in TIC populations in breast cancer cells, this suggests the intriguing possibility that GRP78 may facilitate the acquisition and/or promotion of a stem-like state in breast cancer cell populations. The fact that TIC populations increased dramatically within a short time-frame (e.g. 3 days after GRP78 induction in MCF7 cells) suggests that GRP78 overexpression most likely leads to a proliferative advantage of TIC cells. Nevertheless, it is possible that GRP78 induction can also influence cell fate, and that both mechanisms are contributing to this phenotype. Further experimentation requiring genetic cell fate mapping would be necessary to distinguish between these possibilities. Either way, it is interesting that overexpressing GRP78 leads to both an increase in the number of somatic cells that acquire pluripotency, and an increase in a previously identified breast “cancer stem cell” population [32, 33].

We next examined cell surface functions of GRP78 in in the acquisition and maintenance of pluripotency, and in breast cancer. We found that treatment with an antibody that interferes with cell surface function of GRP78 inhibited both reprogramming efficiency, and pluripotent stem cell and breast cancer cell functions, supporting a cell surface signaling function of sGRP78 in mediating these effects. Further examination of sGRP78 cells within breast cancer revealed that they represented a subset of cells that show a significant increase in genes important in stem cells, and are a subset of a previously identified breast cancer TICs (i.e. a “cancer cell stem cell” population). Strikingly, sGRP78^+^ cells demonstrated a significant increase in *in vitro* and *in vivo* tumorigenicity, suggesting that sGRP78 helps mediate aggressive functions of breast cancer cells.

Previous reports indicate that GRP78 is aberrantly localized to the cell surface in many types of cancer where it has been linked to the regulation of critical signaling pathways [11, 20, 21]. However, as GRP78 is thought to exist majorly as a peripheral protein [34], there have been a number of diverse cell surface binding partners reported for GRP78 that facilitate its activation of downstream signaling cascades, including but not limited to a2M [35], Cripto [20, 22], Par-4 [36] and Kringle 5 [37]. The type of response mediated by sGRP78 is influenced by binding partner, and can also vary depending on the cell type [34]. The preferential surface expression of GRP78 on cancer cells *in vivo* and correlations to poor prognosis and overall metastasis and aggressiveness makes it an attractive chemotherapeutic target. However, without insight into the specific GRP78-dependent mechanisms that are responsible for mediating cancer cell growth and metastasis, it will be difficult to determine how to best target GRP78, or to determine the testing parameters by which to develop the best therapeutic options. Our collective findings support the concept that embryonic mechanisms of GRP78 may be aberrantly retained and/or reactivated in aggressive breast cancer. Ongoing studies from our laboratory are examining the molecular mechanisms by which GRP78 is mediating these effects in both PSCs and cancer. It is our expectation that by focusing on the specific embryonic mechanisms of GRP78 utilized by cancer, this will reveal the specific GRP78-mediated mechanisms that lead to the most aggressive cancer outcomes, and that this information will be critical in focusing future therapeutic targeting of sGRP78 in a clinical setting.

## METHODS

### Cell lines and cell culture

MCF7, MDA-MB-231 and 293T cells (all obtained from ATCC) were grown in Dulbecco’s Modified Eagle’s Medium (DMEM) containing 10% fetal bovine serum (FBS). Human neonatal keratinocytes (Lonza) were grown according to manufacturer’s recommendations. dFib-OCT4^GFP^ fibroblast cells [29] were grown in DMEM containing 10%FBS. Pluripotent stem cells were grown in mTeSR-1 on matrigel as previously described [38, 39]. FiPS4F5 cells [40] and dFib-OCT4^GFP^-iPSC cells [29] have been reported previously. For experiments utilizing doxycycline-inducible treatments, cells were treated with 1µg/ml of doxycycline for the times indicated. In cases requiring GRP78 antibody treatments, GRP78 N-20 antibody (SantaCruz) or relevant goat IgG control (SantaCruz) were used at the concentrations indicated. For both doxycycline and antibody treatments, in cases where multiple days of treatment were required, cells were treated daily with new media containing fresh drug or antibody treatments.

### Plasmid construction and viral production

Retroviral reprogramming plasmids (on pMX backbone) have been described previously [41]. For generation of pMX-GRP78 plasmid, cDNA fragments of GRP78 obtained from pcDNA3-GRP78 [20] were digested with EcoRI, purified and subcloned into EcoRI-linearized pMX plasmid. For viral production, pMX plasmids were cotransfected with packaging plasmids pCMV-VSVG and pCMV-gag-pol-PA as previously described [41].

shRNA (pLVTHM-shScramble and pLVTHM-shGRP78) was generated as previously described using pLVTHM plasmid (shGRP78 sequence: CCATACATTCAAGTTGATA) [29]. Lentivirus was produced by cotransfecting pLVTHM plasmids with packaging plasmids psPAX2 and pMD2.G as previously described [29].

For RFP-GRP78 fusion into pLV-FU-tetO plasmid [29], first tRFP was amplified from pTRIPZ (Addgene) using the forward primer (5’-CACCACCGGTATGAGCGAGCTGATCAAG-3’) and reverse primer (5’-CTCGAGTCTGTGCCCCAGTTTGCT-3’) and the fragment was subcloned into pENTR/D-TOPO using TOPO cloning (pENTR/tRFP). Next, it was digested with XhoI and the fragment of flag-GRP78 [20] was amplified using the forward primer (5’-GGGGCACAGACTCGAAATGAAGCTCTCCCTGGTG-3’) and reverse primer (5’-CCCACCCTTCTCGAGCCTAACAAAAGTT-3’) and subcloned into XhoI-site of pENTR/tRFP with In-Fusion system (pENTR/tRFP-flag-GRP78). The tRFP-flag-GRP78 fragment was amplified with forward primer (5’-GCTTGATATCGAATTCTAACAAAAGTTCCTGAGTCCA-3’) and reverse primer (5’-CCGCGGCCCCGAATTCTAGGCCACCATGAGCGAGCTGATCAAGGAG-3’) from pENTR/tRFP-flag-GRP78 and subcloned into the pLV-FU-tetO vector with In-fusion system. To generate lentivirus pLV-FU-tetO plasmids were cotransfected with packaging plasmids (pMDL, Rev and VSVg) as previously described [29],

### Reprogramming analysis

Keratinocytes (Lonza) or dFib-OCT4^GFP^ were infected with equivalent ratios of retroviruses encoding OSKM (and where indicated with a parallel pMX-GFP control) as previously described [41]. Cells were either replated onto MEFs (Millipore) (keratinocytes) or plated onto Matrigel (dFib-OCT4^GFP^) in their respective media, and then were switched to ES cell medium for iPSC colony formation as previously described [29, 41]. Resulting iPSC colonies were stained for Nanog ∼20 days after infection as previously described [41]. Reprogramming efficiencies were then determined by calculating the number of Nanog positive colonies as a percentage of GFP positive cells.

### RNA isolation and Gene Expression Analysis

RNA was extracted from cell pellets using the RNeasy Mini Kit (Qiagen) according to manufacturer’s recommendations. RNA was reverse transcribed into cDNA using iScript Reverse Transcription Supermix (Bio-Rad), according to the manufacturer’s recommendations. qRT mastermixes were made using SsoAdvanced Universal SYBR Green Supermix (Bio-Rad), according to manufacturer’s recommendations, and qRT was done in a 96-well 7500 Real-Time PCR System (Applied Biosystems). Data was analyzed using Excel and R via the DDCT method standardized to an internal control. Stem cell primers were chosen based on a previously established stem cell array (http://www.sabiosciences.com/rt_pcr_product/HTML/PAHS-405A.html) and are as previously described [29]. Each replicate was compared to its own 18S control, and then compared to its own sorted control population. Following this, the normalized data was averaged across all experiments to determine the mean normalized expression values for each gene in each population. Boxplots and heatmaps were made from the mean normalized or normalized values. All statistical analysis was done using R and plots made using ggplot2.

### Immunofluorescence and immunohistochemistry

Cells were grown on glass coverslips until desired confluency and then were fixed with 4% paraformaldehyde for 15 minutes (min) at room temperature (RT). Cells were permeabilized with cold 0.1% Triton-X/PBS for 15 min at RT and then blocked with cold 2% FBS/PBS for 30 min at RT. Primary antibody was diluted into 2% FBS/PBS and cells were incubated O/N at 4C. Following washing cells were incubated with secondary antibody in 2% FBS/PBS for 2 hours at RT. Finally, the cells were incubated with DAPI diluted in PBS for 10 min at RT and washed before being mounted onto slides with Vectashield Hard Set (Vector Laboratories). Single-planed images were taken on a Nikon C2 confocal microscope using a 40X oil-immersion lens, and figures were arranged using Adobe Photoshop or Illustrator. All fluorescent images shown within a figure were acquired with the same exposure time.

### MTT Assay

Cells were plated in a 96-well plate before being treated daily with media only, or media containing doxycycline and/or GRP78 N-20 antibody or IgG control. The cells were next treated with cisplatin at the concentrations and times indicated. Following incubation, the media was aspirated from the cells and replaced with 50uL of fresh media containing 0.5mg/mL Thiazolyl Blue Tetrazolium Bromide (MTT) reagent (Sigma) per well. The cells were incubated for 4 hours at 37 degrees. After incubation, 150 uL of DMSO was added to each well and mixed to completely dissolve the solution. Absorbance was measured in plate reader at 570nm.

### Flow Cytometry and FACS

MCF7 cells were grown on 10cm plates until complete confluency. Cells were harvested (using EDTA/PBS; Invitrogen) and aliquoted into individual samples containing 1×10^6^ cells for labeling. Samples were stained with either anti-rabbit GRP78 ET-21 (Sigma) antibody, or CD24-APC or CD44-FITC (eBioscience) for 1 hr at 4°C. Primary antibody (GRP78 ET-21) was used at 5µg/1million cells. When appropriate, cells were incubated with anti-rabbit 488 secondary antibody (Invitrogen) for 1 hr at 4°C. Cells were then either analyzed via flow cytometry (BD Fortessa) or sorted (BD FACSAria Cell Sorter) based on the staining of both a secondary-only and IgG (Sigma) control (for GRP78-labeled cells) or single-stained controls (APC/FITC). When cells were sorted for the *in vivo* experiments, all cells were labeled with a Dil Stain (1,1’-Dioctadecyl-3,3,3’,3’-Tetramethylindocarbocyanine Perchlorate) (Invitrogen) immediately following sorting, just before injection, according to the manufacturer’s recommendations.

### Soft Agar Colony Forming Assay

Sorted MCF7 populations (as described) were plated at 30,000 cells per well in 1.5 ml of growth media plus 0.4% low-melt agarose (Fisher Scientific) and layered onto a 3 ml bed of growth media with 0.5% low-melt agarose (Fisher Scientific). Cells were fed daily with 1 ml of growth media. At the indicated time, growth media was removed and viable colonies were stained using Iodonitrotetrazolium chloride (INT-Violet) (Sigma). Colony number and size was determined using ImageJ analysis (Bethesda, MD, USA).

### *In Vivo* Experiments

All animal experiments were performed ethically and in accordance with IACUC protocol approved by the University of Notre Dame IACUC committee. Rag1−/− (C.129S7(B6)-Rag1^tm1Mom^/J) mice were purchased from The Jackson Laboratory (Bar Harbor, ME). All mice were eight weeks or older prior to experimental procedures. *In vivo* tumor seeding and growths were formed by injection of 20,000 cells (DiI-labeled MCF7-total; DiI-labeled sGRP78^+^; or DiI-labeled-CSCs (isolated and labeled as described above) suspended in 150uL serum-free RPMI media into the left cardiac ventricle. Prior to and during injection of cancer cells, mice were anesthetized with isoflurane. Mice were sacrificed after two days post-injection for short term experiments and four weeks post-injection for long term experiments. Organs were immediately extracted and fixed in 4% paraformaldehyde overnight, and then washed and stored in PBS. The presence of fluorescent cells was manually counted for lungs and brains as indicated. For confocal analysis, small pieces of lung tissue were cut off from the collected lungs, stained with DAPI and imaged by confocal microscopy to examine DiI stained cells.

### Statistical Analysis

Results are shown as mean values ± standard error of the mean (SEM). Remaining statistics were performed using unpaired two-tailed Student’s *t*-tests, or Anova when doing multiple comparisons. *p* values<0.05 were considered statistically significant.

## Supporting information

Conner et al_Supplementary Figures

## ACKNOWLEDGMENTS

This work was supported in part by a Walther Cancer Foundation Advancing Basic Cancer grant, and by grant UL1TR00110 from the Indiana Clinical and Translational Sciences Institute (to ADP). CC was supported in part by grant TL1TR001107 from the National Institutes of Health, National Center for Advancing Translational Sciences, Clinical and Translational Sciences Award. AEY was supported in part through The University of Notre Dame College of Science Deans’ Fellowship, and RCG through a Hiller Family Research Fellowship. We thank the Gallagher Family for their generous support of stem cell research at the University of Notre Dame.

## AUTHORSHIP CONTRIBUTIONS

CC, TWL, AEY, RCG, MZW performed experiments and analysis. IHG, SY contributed to *in vivo* experiments. CC and TWL contributed to writing of the manuscript. SR, YH, TH provided plasmid constructs. JCIB, PCG, JAK discussed project findings, and PCG, JAK provided reagents and relevant expertise on project design. ADP designed and supervised the research project, analyzed the data, and wrote the manuscript. All authors reviewed the manuscript.

## DATA AVAILABILITY

All data generated or analyzed during this study are included in this published article (and its Supplementary Information files).

