## Supplementary material for "GRP78 promotes stemness in normal and neoplastic cells": Conner et al_Supplementary Figures

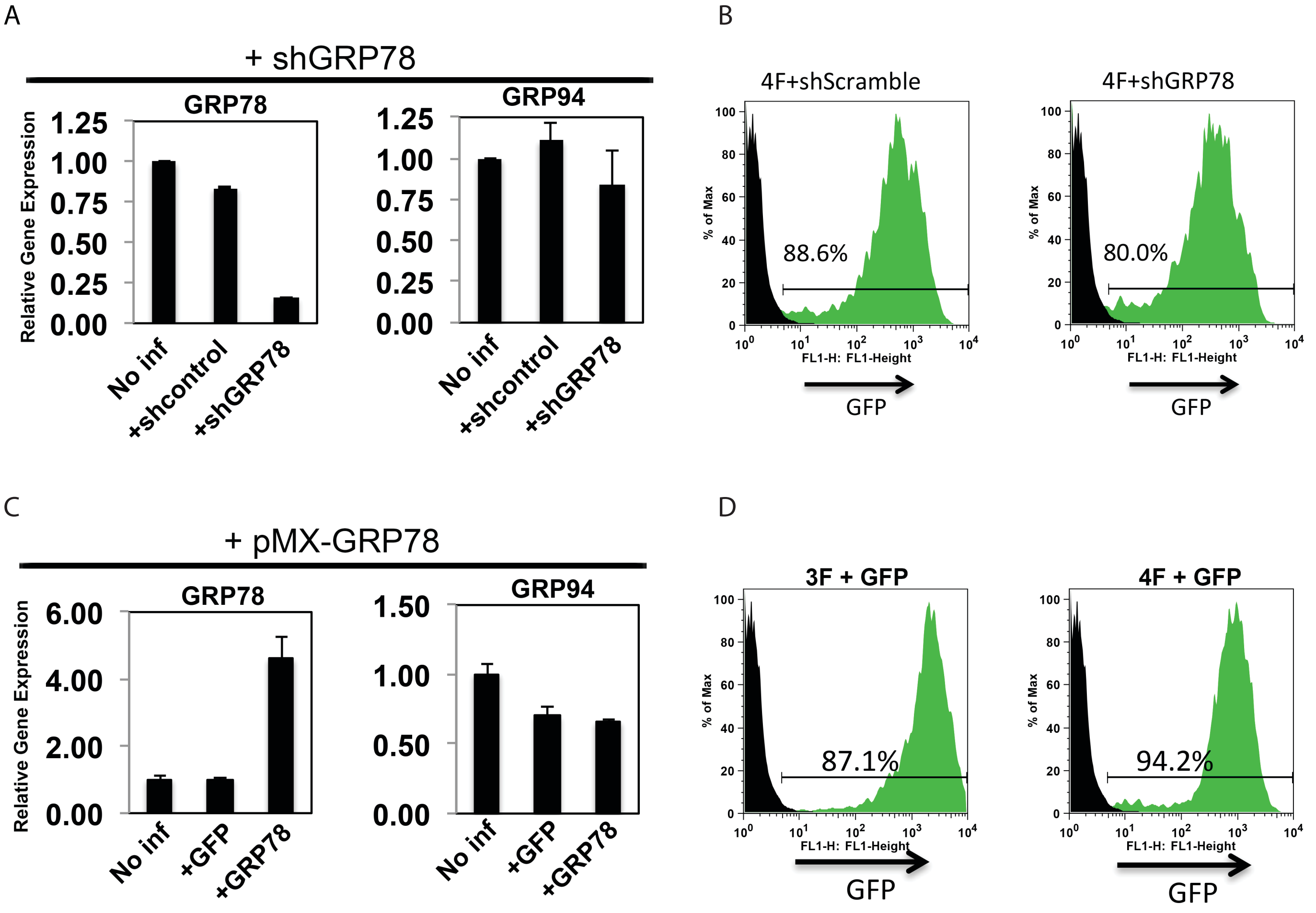


**Figure S1. Validation of GRP78 knockdown and overexpression systems used in reprogramming experiments (related to Figure 1).** (A) Gene expression analysis of GRP78 or GRP94 (another heat shock family member) in human keratinocytes of shGRP78 or a shScramble control, showing that shGRP78 selectively knocks down GRP78 transcript levels. (B) Flow cytometry analysis of keratinocytes infected with shGRP78 or shScramble lentiviruses containing GFP, relative to uninfected controls, show similar infection efficiencies for both. (C) Gene expression analysis of GRP78 or GRP94 in keratinocytes following infection with pMX-GRP78 (to overexpress GRP78) or a pMX-GFP control relative to uninfected controls, confirming that GRP78 is selectively overexpressed following pMX-GRP78 infection. (D) Flow cytometry analysis of keratinocytes infected with pMX-GRP78 or pMX-GFP retroviruses relative to uninfected controls, show similar infection efficiencies for both. [Note that equivalent numbers of GFP positive cells were used when performing reprogramming efficiencies experiments described in Figure 1].

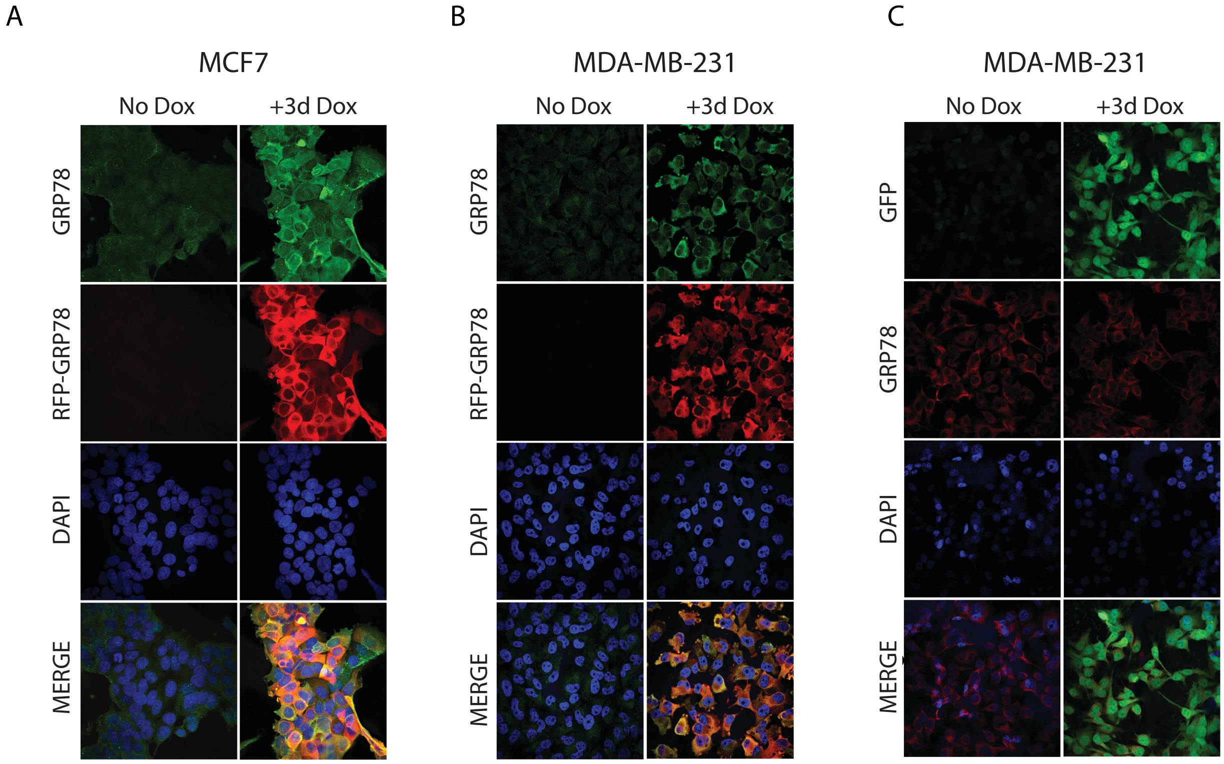


**Figure S2. Inducible RFP-GRP78 fusion in breast cancer cells colocalizes with GRP78 (related to Figures 2 and 3)**. (A-B) MCF7-RFP-GRP78 (A) and MDA-MB-231-RFP-GRP78 (B) cells were treated with doxycycline for 3 days (+3d Dox) or were left untreated (No Dox). Cells were fixed and stained with a GRP78 antibody and 4,6-Diamidino-2-phenylindole (DAPI) and imaged. GRP78 (green) appears to be increased in the presence of doxycycline and was consistently found to be colocalized with RFP-GRP78. (C) An inducible GFP control with the same vector backbone as RFP-GRP78 shows that doxycycline alone was not enough to induce the overexpression of GRP78 (red).

A

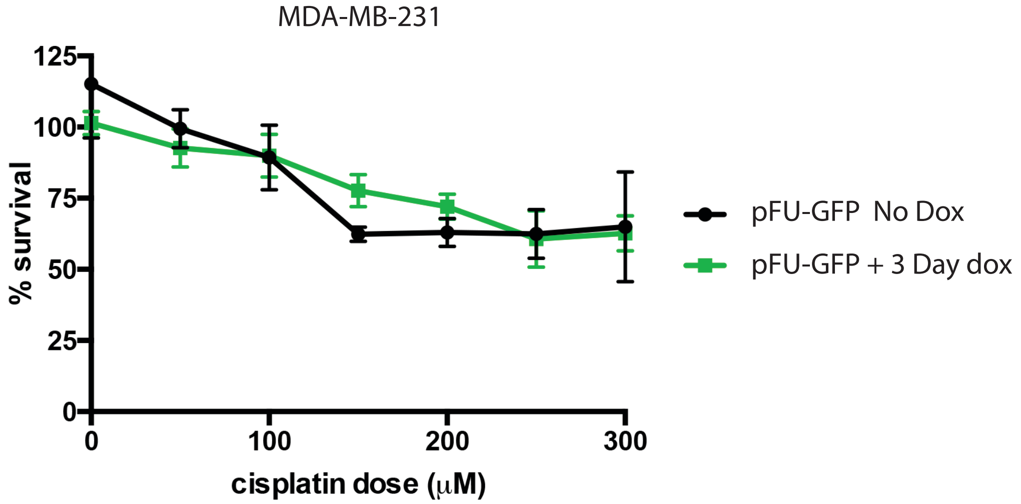


B

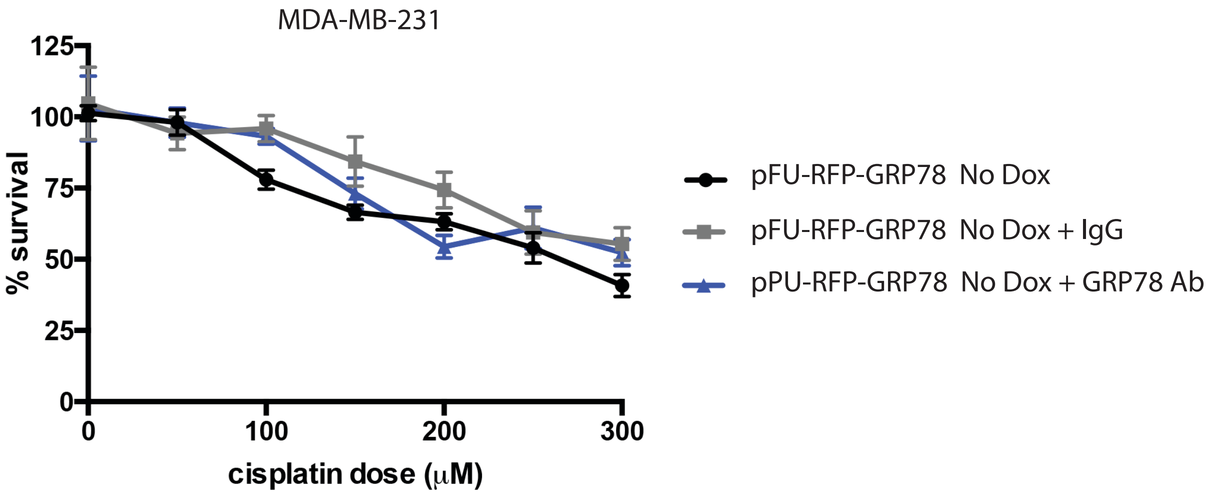


**Figure S3. Confirmation that cisplatin resistance in MDA-MB-231 cells is mediated by overexpression of GRP78 (related to Figure 2).** (A) MDA-MB-231 breast cancer cells that overexpressed GFP under a doxycycline (Dox)-inducible promoter were used in an MTT-based survival assays, and show no resistance to cisplatin-induced apoptosis. (B) In the absence of doxycycline treatment, MDA-MB-231 cells that contain inducible RFPGRP78 show no resistance to cisplatin-induced apoptosis, and no response to treatment with a GRP78 inhibitory antibody.

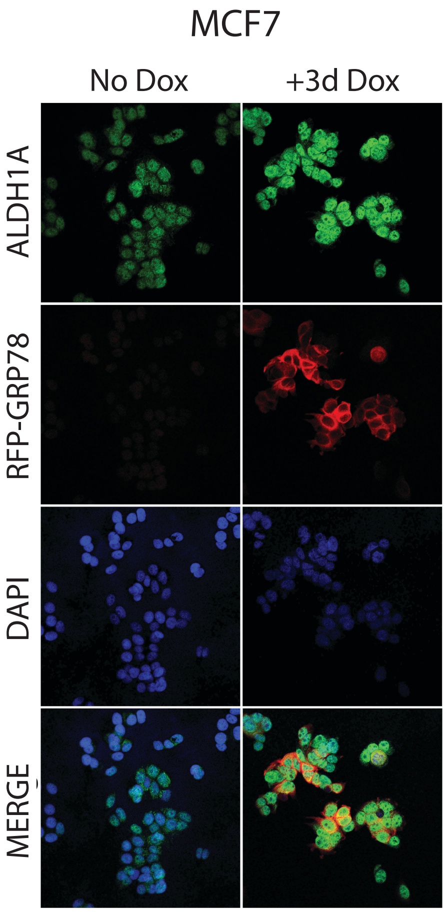


**Figure S4: ALDH1A is increased in MCF7 cells following GRP78 overexpression (related to Figure 3).** MCF7-RFP-GRP78 cells were treated with doxycycline for 3 days (+3d Dox) or were left untreated (No Dox) and then fixed and stained with an ALDH1A antibody and 4,6-Diamidino-2-phenylindole (DAPI). ALDH1A (green) shows an increase in expression in cells that overexpress GRP78 (red).
